# Integration of Mass Cytometry and Mass Spectrometry Imaging for Spatially Resolved Single Cell Metabolic Profiling

**DOI:** 10.1101/2023.08.29.555282

**Authors:** Joana B Nunes, Marieke E Ijsselsteijn, Tamim Abdelaal, Rick Ursem, Manon van der Ploeg, Bart Everts, Ahmed Mahfouz, Bram Heijs, Noel FCC de Miranda

**Affiliations:** Pathology, Leiden University Medical Center, Leiden, The Netherlands; Department of Radiology, Leiden University Medical Center, Leiden, The Netherlands; Systems and Biomedical Engineering Department, Faculty of Engineering Cairo University, Giza, Egypt; Pattern Recognition and Bioinformatics, Delft University of Technology, Delft, The Netherlands; Center for Proteomics and Metabolomics, Leiden University Medical Center, Leiden, The Netherlands; Parasitology, Leiden University Medical Center, Leiden, The Netherlands; Human Genetics, Leiden University Medical Center, Leiden, The Netherlands; The Novo Nordisk Foundation Center for Stem Cell Medicine (reNEW), Leiden University Medical Center, Leiden, the Netherlands

## Abstract

Integration of spatial omics technologies can provide important insights into the biology of tissues. We combined mass spectrometry imaging-based metabolomics and imaging mass cytometry-based immunophenotyping on the same single tissue section to reveal metabolic heterogeneity within tissues and its association with specific cell populations like cancer cells or immune cells. This approach has the potential to greatly increase our understanding of tissue-level interplay between metabolic processes and their cellular components.

## Main text

Metabolism is an essential aspect of biological systems that must be considered to comprehend tissue homeostasis and pathogenesis (1, 2). Single-cell proteomic and transcriptomic analyses have significantly advanced our understanding of metabolic variations across different cell populations, particularly immune cells, to uncover metabolic heterogeneity as well as providing insights into the intricate links between cell phenotypes and metabolic profiles (3-5). However, these approaches extract cells from their natural context, lacking spatially-resolved information and disregarding cellular interactions. Furthermore, relying on surrogate markers, such as enzymes or their transcripts, to infer metabolic states, overlooks the direct detection of metabolite abundance, which remains the most reliable approach.

We developed a novel multimodal imaging approach for the integrated analysis of metabolites and immunophenotypes in human tissues. This was achieved by integrating the experimental workflows and the data generated from spatial metabolomics using matrix-assisted laser desorption/ionization mass spectrometry imaging (MALDI-MSI) and spatial immunophenotyping by imaging mass cytometry (IMC). We developed and optimized a wet-lab protocol that allows the application of both technologies on the same tissue section. Additionally, we have implemented a data integration strategy that enables the relative quantification of metabolites (measured by MALDI-MSI) at single cell level (defined by IMC) (Figure 1A).

**Figure 1.**
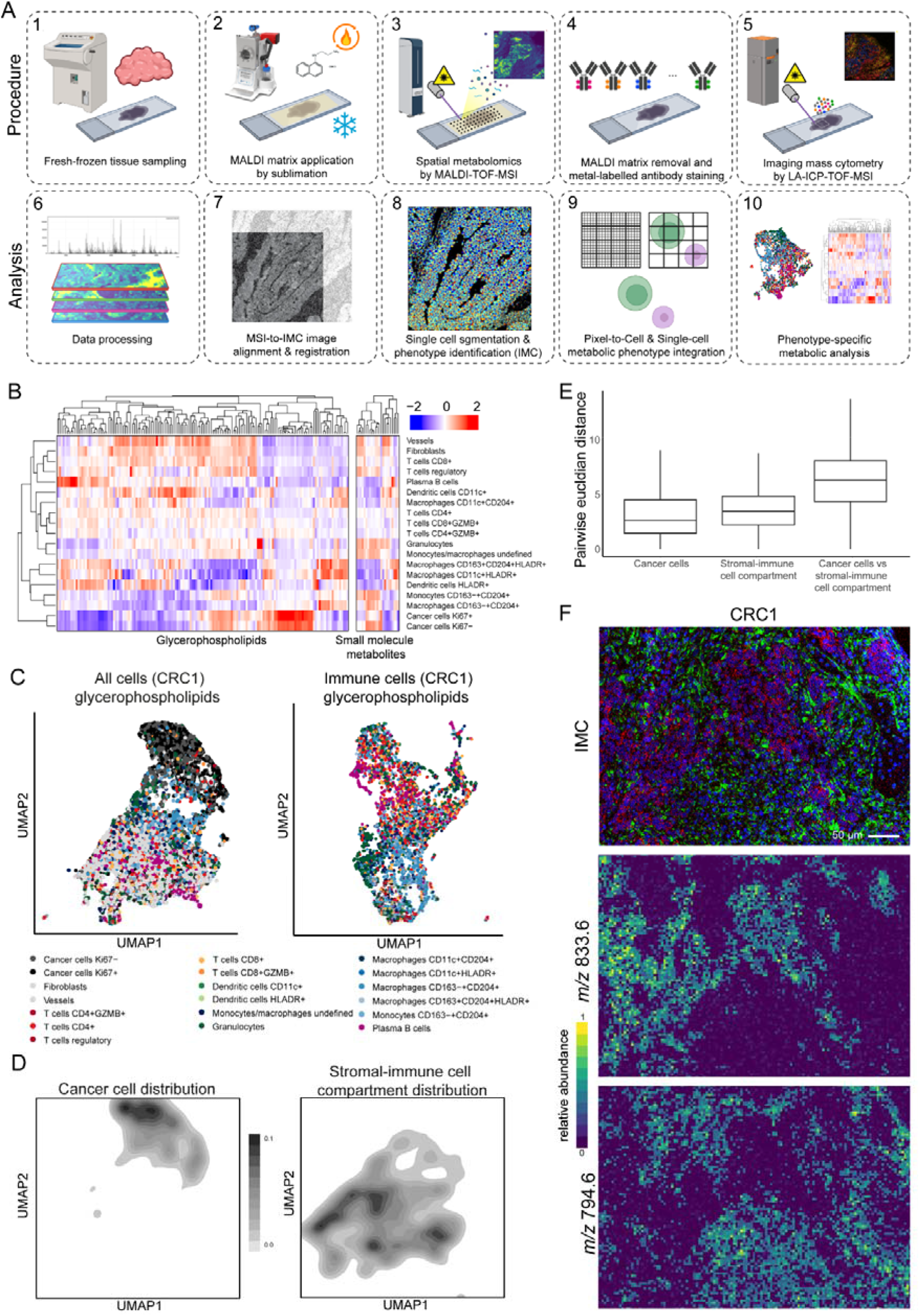
A. Integrated workflow of consecutive MALDI-MSI and IMC analyses. Fresh-frozen tissue sections were cut (1) and treated with MALDI matrix (2). MALDI-TOF-MSI was performed to obtain spatial metabolomics data (3), followed by matrix removal (4). Sections were then labelled with metal-conjugated antibodies and IMC was used to analyze the areas previously imaged by MALDI-TOF-MSI (5). Both datasets were pre-processed (6) and co-registered using visual landmarks (7). Cells segmentation and phenotype identification were performed (8) and MALDI-MSI-derived metabolic abundances were assigned to each cell (9), enabling downstream analysis (10). B. Scaled metabolite expression profiles across cell populations identified in CRC1. Hierarchical clustering was guided by glycerophospholipid abundances (heatmaps of CRC2 and CRC3 are shown in supplementary figure 2D). C. UMAP embedding of all cells (16952 cells, left panel) or immune cells (6523 cells, right panel) of CRC1, clustered by glycerophospholipids (*m/z* >400). Phenotypes of labelled cells are visualized on the UMAP embedding. D. distribution of cancer cell phenotypes and immune/stromal phenotypes corresponding to the UMAP embedding of C (left). E. Pairwise distances derived from the UMAP embedding between cancer cells, cancer cells and the stromal-immune cell compartment or the stromal-immune cell compartment only, visualized in a boxplot with median distance and interquartile range. F. Top: representative IMC image of CRC1 highlighting cancer cells and stromal cells (keratin in red, vimentin in green and DNA in blue). Bottom: MALDI-MSI image of the same region of interest (ROI) showing differentially abundant metabolites in cancer cells (*m/z* 833.6) and the stromal-immune cell compartment (*m/z* 794.6) displayed using viridis colors with saturated pixels above the 99^th^ percentile.

To achieve single cell resolution, we performed MALDI-MSI and IMC at spatial resolutions of 5×5 μm and 1×1 μm pixel areas, respectively, on fresh-frozen colorectal cancer tissue sections. MALDI-MSI was conducted prior to IMC due to the tissue-ablative nature of IMC. Laser parameters for MALDI-MSI were carefully adjusted to minimize tissue damage while ensuring ionization (Supplementary Figure 1A). Typically, MALDI-MSI is performed on 10-12 μm thick tissue sections, while IMC is applied on 4-5 μm thick sections to minimize cell overlap. We evaluated the quality of MALDI-MSI measurements on 5 μm sections and found no discernable differences in signal intensity or spatial distributions of metabolic features when compared to 10 μm sections (Supplementary Figure 1B). Based on these results, we decided to employ 5 μm sections for subsequent analyses.

Following the acquisition of MALDI-MSI data and the removal of excess MALDI matrix, we evaluated the impact of this procedure on IMC-based antibody-mediated antigen detection. We initially used an IMC panel designed for snap-frozen tissues (6) but encountered challenges such as low signal intensity from antibody-derived metal ions and the failure of several antibodies crucial for morphological evaluation of tissues (Supplementary Figure 1C). We hypothesized that the room temperature processing of slides during the MALDI-MSI procedure might compromise tissue stability and epitope integrity. To address this issue, we examined an adapted version of an antibody panel developed for formalin-fixed paraffin embedded (FFPE) tissues (Supplementary Table 1) (7). Additionally, we incorporated a formalin fixation step following the removal of the MALDI matrix. Remarkably, this modified approach resulted in improved metal ion signal intensity and the specificity of antigen detection compared to using the antibody panel originally developed for snap-frozen tissues (Supplementary Figure 1C).

We applied our optimized protocol on three tumor samples, including one mismatch repair (MMR) deficient colorectal cancer (CRC1) and two MMR proficient colorectal cancers (CRC2 and CRC3). To merge the output data from MALDI-MSI and IMC, we performed image co-registration based on visual features that contained recognizable landmarks such as empty areas or epithelial structures (Supplementary Figure 2A). However, due to the differences in resolution and pixel size between MALDI-MSI and IMC, we needed to account for these variations to define metabolite abundance at the single-cell level. First, we attributed the metabolite abundance of each MALDI-MSI pixel to the corresponding 25 overlapping IMC pixels (Online methods, Supplementary figure 2B). Next, we identified cells through cell segmentation using the DNA, keratin (epithelial cells), and vimentin (stromal cells) images from the IMC data (8). Subsequently, using the previously assigned pixel metabolite abundance, we calculated the relative metabolite abundance per cell. Additionally, by leveraging the cell marker expression from the IMC data, we identified a total of 19 cellular phenotypes, including cancer cells, macrophages, and T cells (Supplementary Figure 2C). As MSI pixels are 25 μm^2^, multiple cells can correspond to a single MSI pixel, which can result in falsely assigned metabolite profiles. But we observed that the majority of MSI pixels contained zero or one cell and thus we reasoned that this would not substantially influence downstream analyses (Supplementary Figure 2D).

We performed hierarchical clustering analysis to evaluate metabolite abundance across all 19 identified cell populations. The analysis revealed distinct clusters of glycerophospholipids associated with various cell types, particularly cancer cells, plasma B cells and macrophages (Figure 1B and Supplementary Figure 2E). To investigate whether different metabolic profiles could be distinguished within the same cell subsets, we focused our analysis on CRC1, which exhibited high immune infiltration, in line with its MMR-deficient status. We conducted UMAP dimensionality reduction on either all cells combined or immune cells only using the metabolite abundances as features. Moreover, as glycerophospholipids are expected to be more differentially abundant between cell types compared to small molecule metabolites, both metabolite types were analyzed separately. Consistent with the distinct metabolite profile observed in Figure 1B, cancer cells localized separately from stromal and immune cells in the UMAP embedding (Figure 1C, D and Supplementary Figure 3A). This separation was confirmed by a higher pairwise distance between the cancer cell compartment and the stromal-immune cell compartment compared to within these subsets (Figure 1E).

To identify metabolites with different abundances between cancer and stromal-immune cell compartments, we performed differential feature analysis (Supplementary Figure 3B). Two of the top differentiating metabolites between these tissue compartments can be visualized on Figure 1F. These metabolites were found to overlap with Keratin (for cancer cells) or Vimentin (for the stromal-immune cell compartment), validating the robustness of the MALDI-MSI IMC combination approach (Figure 1F). Although cancer cells and the stromal-immune cell compartment exhibited separated clustering, immune cells only partially grouped by cell type in the UMAP embedding (Figure 1C), indicating differences in metabolite abundances within the same cell populations. To confirm that little clustering by cell type occurs, we conducted k-means clustering of the immune cells based on glycerophospholipid features and compared the results with the previously assigned cell types using a confusion matrix (Supplementary Figure 3C and D). The analysis demonstrated no clear association between the IMC and glycerophospholipid clusters, confirming the presence of metabolite variations within the same cell populations. This phenomenon was particularly evident in macrophages, which formed clusters but also scattered throughout the embedding (Figure 1C and Supplementary Figure 3E). Therefore, we conducted further investigations to explore these observations within the macrophage population.

To investigate the distinct metabolic profiles of macrophages/monocytes, we conducted differential feature analysis, comparing them with all other cells in the dataset. This analysis revealed several metabolites with increased abundance specifically in macrophages/monocytes. (Figure 2A and B). We then visualized these cells in a UMAP embedding based on metabolite profiles and observed different clusters of cells that were not clearly associated with assigned cell phenotypes (Figure 2C). In contrast, k-means clustering of the macrophages and monocytes demonstrated significant differences in glycerophospholipid abundance between the clusters (Figure 2D and E), providing further support for the hypothesis that different metabolic profiles exist within the macrophage/monocyte population. Overall, our multimodal imaging and analysis approach employed in this study enables the evaluation of metabolites at the single-cell resolution and emphasizes the presence of differences in metabolite abundance both between and within cellular phenotypes.

**Figure 2.**
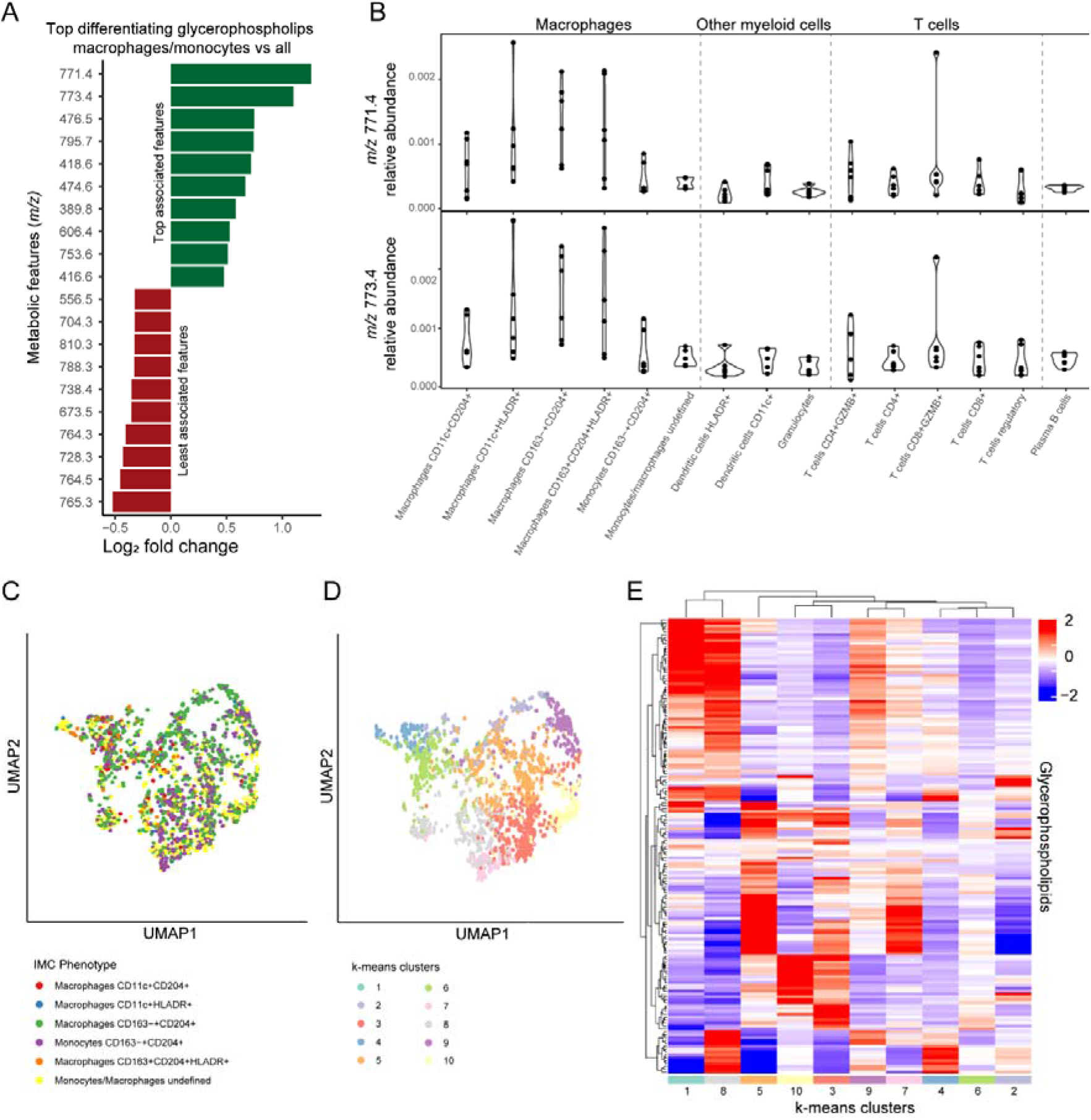
A. Differential glycerophospholipid features distinguishing macrophages/monocytes from other cells, calculated for all images in the dataset. B. Violin plots showing the relative abundance per cell of the two most differentially abundant metabolites in macrophages/monocytes. Each dot represents an image (*m/z* 771.4 assigned to PI(30:5) [M-H]^-^ and *m/z* 773.4 assigned to PI(30:4) [M-H]^-^). C. UMAP embedding of macrophages/monocytes in CRC1, utilizing glycerophospholipids as features (*m/z* >400) as features. Cells are labelled by IMC phenotypes D. k-means clusters of macrophages/monocytes based on glycerophospholipids, visualized on the UMAP embedding (2C). E. Relative abundance of glycerophospholipids in the k-means clusters, visualized in Figure 2D.

The field of cancer metabolism faces a significant challenge in assessing the metabolic profile of immune cells within the complex microenvironment of tumors. Previous studies have attempted to address this challenge by combining MALDI-MSI with immunofluorescence (IF) imaging or IMC on consecutive tissue sections. However, these approaches are difficult to integrate and do not provide true single-cell metabolic profiles (9, 10). Other studies have performed IHC or IF on the same section after MALDI-MSI, but they are limited in the number of targets that can be evaluated (11, 12). Although DESI-MSI and IMC have been successfully combined on the same tissue section, the compromised spatial resolution of the DESI-MSI platform prevents achieving single-cell resolution (13). Furthermore, the development of MALDI-IHC, which analyzes photocleavable peptide-tagged antibodies by MALDI-MSI, allows a similar approach but with compromised spatial resolution for immunophenotyping compared to IMC (14, 15). In this study, we developed and demonstrated an optimized workflow that combines the detection of metabolites using MALDI-MSI with the identification of cellular phenotypes through IMC on a single tissue section, thereby overcoming previously posed challenges. Moreover, while we focused on human colorectal cancer tissue in this study, our methodology is suited for the analysis of diverse tissue types.

In conclusion, our methodology provides a powerful tool for advancing the characterization of tissues by simultaneously uncovering their cellular composition and the metabolic profiles of cells. This capability holds immense importance, for instance in the field of cancer immunology, where cellular metabolism plays a critical role in shaping immune cell function and in the efficacy of immunotherapeutic interventions. By applying our approach to cancer samples from patients undergoing immunotherapy, we anticipate uncovering valuable metabolism-associated features that can be correlated with treatment response.

## Acknowledgements

Noel FCC de Miranda is funded by the European Research Council (ERC) under the European Union’s Horizon 2020 Research and Innovation Programme (grant agreement no. 852832). Bram Heijs is funded by the Dutch Kidney Foundation Kolff+ Jr. talent (grant no. 21OK+015) and the Novo Nordisk Foundation Center for Stem Cell Medicine (reNEW),supported by Novo Nordisk Foundation grants(NNF21CC0073729). Joana B Nunes is funded by the Leiden university medical centre (LUMC) via the internal regulation for MSCA-IF Seal-of-Excellence. Bart Everts is funded by an LUMC fellowship. The authors thank Paola Tasca for technical assistance.

## Data availability

Raw IMC and MSI data as well as combined datasets are available via Figshare (https://figshare.com/s/c58ddc70fa8dc0602842).

## Code availability

Scripts and command lines used to analyse the presented data are available at Figshare (https://figshare.com/s/c58ddc70fa8dc0602842).

## Methods

### Sample collection/Sampling

Material was used of three CRCs from patients that have given informed consent under the study protocol P15.282, approved by the Medical Ethical Committee of the Leiden University Medical Centre. Patient samples were anonymized and handled according to the medical ethical guidelines described in the Code of Conduct for Proper Secondary Use of Human Tissue of the Dutch Federation of Biomedical Scientific Societies. After surgery, cancer tissue was snap-frozen and stored at -80 degrees.

Using a micro cryostat (CM3050 S, Leica), 5 μm-thick tissue sections were thaw-mounted on indiumtin-oxide (ITO)-coated microscope slides (Bruker Daltonics GmbH), and immediately processed for MALDI-MSI analysis.

### Matrix-assisted laser desorption/ionization mass spectrometry imaging (MALDI-MSI)

Samples from the cryomicrotome were transferred on dry ice and equilibrated to room temperature using vacuum freeze drier (Alpha 2-4 LSCbasic, Christ, Osterode am Harz, Germany) for 20 minutes. Following, the MALDI matrix, 1-napthylyl ethylenediamine dihydrochloride (NEDC), was sublimated onto the slide using a Sublimator T1 (HTX Imaging). Sublimation was performed at 180°C for 6 minutes and which resulted in a matrix coverage of 31.73 pmol/mm^2^. MSI analysis was performed on a rapifleX MALDI-TOF/TOF-MS platform (Bruker Daltonics GmbH). Single-pixel mass spectra were recorded at a 5×5 μm^2^ spatial resolution covering a *m/z* range between 60-1,400 Th, and summing 50 laser shots. The flexImaging software (v5.0, Bruker Daltonics GmbH) was used to define measurement regions of approximately 1×1 mm^2^. Post measurement, samples were immediate processed for the imaging mass cytometry analysis. Raw data files were loaded into the SCiLS Lab software (v2023b, Bruker Daltonics GmbH), single-pixel spectra were baseline corrected using a TopHat algorithm (window 200) and per measurement region an imzML was exported. imzML files were loaded and further processed using rMSI and rMSIproc (16, 17). Spectral processing included per pixel spectral alignment (maximum shift, 25 ppm; iterations, 1), peak picking (signal-to-noise ratio ≥ 15), recalibration against a list of known *m/z* features, and peak binning (datapoints per peak, 8 datapoints). Peaklist filtering was performed by discarding peaks outside the expected mass defect windows for i) small molecule metabolites below *m/z* ≤ 400 Th based on the Human Metabolome Database (https://HMDB.ca), and ii) glycerophospholipids above *m/z* > 400 Th (lipid classes: glycerophospholipids [GP], fatty acids [FA], glycosphingolipids [SP]) based on LIPIDMAPS (https://lipidmaps.org). Remaining peaks were manually curated, and peaks without clear spatial distribution were discarded. The final peak matrix was exported as *m/z*-specific .tiff files.

### Imaging mass cytometry (IMC)

After MALDI-MSI data acquisition, excess MALDI matrix was removed by washing the tissue slides with 100% ethanol, ice cold acetone, 100% ethanol and 70% ethanol for 5 minutes each, followed by a 2-minute wash with 50% ethanol and 3× 1-minute washes with 1× PBS. The tissues were then fixed with 10% formalin for 1 hour at room temperature. Antigen retrieval, blocking and labeling with metal-conjugated antibodies (Supplementary Table 1) was performed as previously described by IJsselsteijn *et al* (7). To obtain the data in Supplementary Figure 1B, tissue fixation was performed after excess MALDI matrix removal with 4% paraformaldehyde for 30 minutes at 4°C and the IMC staining according to the protocol of Guo *et al*. (6). Data from 1,100×1,100 μm^2^ areas was acquired using a Hyperion™ Mass Cytometry System (Standard BioTools), exported as .mcd files and visualized using MCD™ Viewer (1.0.5, Standard BioTools).

Phenotype identification and counting was done as described previously (8). In short, .mcd files were converted to TIFF images and normalized by saturating pixels above 99^th^ percentile. Next, background removal was performed using the random forest classifier of Ilastik after which all remaining pixels were binarised to 1, with background set to 0. Cell segmentation masks were created using Ilastik and CellProfiler using the keratin, vimentin/CD45 and DNA images and marker intensities per cell were extracted using ImaCytE. Cells were analyzed by t-SNE in Cytosplore and by mean-shift clustering, cells forming visual neighborhoods were grouped to define cellular phenotypes. R-studio (2022.07.1, R version 4.2.0) was used for counting of phenotypes per sample and visual representation.

### MSI and IMC data integration

For each tumor sample, MALD-MSI and IMC data were combined in python, by co-registering the two image sets based on visual features in both datasets showing clear landmarks, like empty areas or epithelial structures. Excess pixels, that can be found on the borders of the images belonging only to one dataset, are excluded, retaining only co-registered pixels. Since MALDI-MSI resolution is 5×5 μm^2^ and IMC resolution is 1×1 μm^2^, each MALDI-MSI pixel maps to 25 IMC pixels.

Let *X*_*i*_ = {*x*_*1*_,*x*_*2*_,…, *x*_*N*_} be the set of IMC pixels where *N* is the total number of co-registered pixels, and *Y*_*j*_ = {*y*_1_,*y*_2_,…, *y*_*M*_} be the set of MALDI-MSI pixels where *M* is the total number of co-registered pixels, such that *M* = *N*/25. Each maps to 25 *x*_*i*_’s, while *x*_*i*_ maps to (is part of) a single, therefore *y*_*j*_ *= f(x*_*i*_ ) where *f()* is a defined mapping function returning the corresponding *y*_*j*_ pixel for the input *x*_*i*_ pixel.

For both modalities, each pixel represents a 1D vector containing the features measured by that modality. Cells were segmented using the high-resolution IMC dataset, let *c*_*k*_= {*c* _*1*_,*c*_*2*_,…, *c*_*k*_} be the set of segmented cells and *n* is the total number of cells. Each cell *c*_*k*_ is a set of several IMC pixels *x c*_*i*_’s, while one IMC pixel *x*_*i*_ pixel is part of only one cell. To obtain the cellular IMC expression data, we define

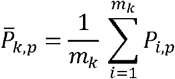

Where 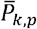 is the cellular expression of IMC protein feature *p* in cell *c*_*k*_, *p*_*i,p*_ is the pixel expression of protein feature *p* in IMC pixel *x* _*i*_, and *m*_*k*_ is the number of IMC pixels corresponding to cell *c*_*k*_. In other words, the cellular expression of the IMC data is the average expression of the IMC pixels corresponding to each cell. Similarly, let *T* _*j,k*_ be the pixel expression of metabolic feature *t* in pixel *y*_*j*_, therefore:

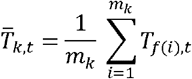

Where 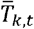 is the cellular expression of metabolic feature *t* in cell *c*_*k*_, *T*_*f(i),t*_ is the pixel expression of metabolic feature *t* in MALDI-MSI pixel mapping to IMC pixel *x*_*i*_. In other words, the cellular expression of the MALDI-MSI data is the weighted-average expression of the MALDI-MSI pixels corresponding to each cell, where each MALDI-MSI pixel is weighted by how many IMC pixels are mapping to it. It is important to note that one MALDI-MSI pixel *y*_*i*_ may contribute to the expression of one or more cells.

### Data analysis

Cellular phenotypes and metabolite expression per cell were combined in R-studio and metabolite expression counts were normalized to relative expression per cell. R-studio was used for all consecutive analyses. For each sample, data from two images were combined and mean relative metabolite expression was determined for each cellular phenotype and visualized in a heatmap per sample using the ComplexHeatmap R package (version 2.14.0). Differential feature expression analysis was done by Wilcoxon rank test on the group of interest versus the remaining cells in the dataset and visualizing the features by fold-change and P-value. Finally, for dimensionality reduction, the cells were visualized in UMAP (Uniform Manifold Approximation and Projection) using the umap R package (version 0.2.10.0) using the following parameters: n_neighbors of 5, min_dist of 0.005 and n_epochs of 1,000 and cell phenotypes or k-means clusters were visualized on the embedding.

## References

1. Wang G, Heijs B, Kostidis S, Rietjens RGJ, Koning M, Yuan L, et al. Spatial dynamic metabolomics identifies metabolic cell fate trajectories in human kidney differentiation. Cell Stem Cell. 2022;29(11):1580–93 e7.

2. Pavlova NN, Zhu J, Thompson CB. The hallmarks of cancer metabolism: Still emerging. Cell Metab. 2022;34(3):355–77.

3. Hrovatin K, Fischer DS, Theis FJ. Toward modeling metabolic state from single-cell transcriptomics. Mol Metab. 2022;57:101396.

4. Artyomov MN, Van den Bossche J. Immunometabolism in the Single-Cell Era. Cell Metab. 2020;32(5):710–25.

5. Purohit V, Wagner A, Yosef N, Kuchroo VK. Systems-based approaches to study immunometabolism. Cell Mol Immunol. 2022;19(3):409–20.

6. Guo N, van Unen V, Ijsselsteijn ME, Ouboter LF, van der Meulen AE, Chuva de Sousa Lopes SM, et al. A 34-Marker Panel for Imaging Mass Cytometric Analysis of Human Snap-Frozen Tissue. Front Immunol. 2020;11:1466.

7. Ijsselsteijn ME, van der Breggen R, Farina Sarasqueta A, Koning F, de Miranda N. A 40-Marker Panel for High Dimensional Characterization of Cancer Immune Microenvironments by Imaging Mass Cytometry. Front Immunol. 2019;10:2534.

8. Ijsselsteijn ME, Somarakis A, Lelieveldt BPF, Hollt T, de Miranda N. Semi-automated background removal limits data loss and normalizes imaging mass cytometry data. Cytometry A. 2021.

9. Tuck M, Grelard F, Blanc L, Desbenoit N. MALDI-MSI Towards Multimodal Imaging: Challenges and Perspectives. Front Chem. 2022;10:904688.

10. Goossens P, Lu C, Cao J, Gijbels MJ, Karel JMH, Wijnands E, et al. Integrating multiplex immunofluorescent and mass spectrometry imaging to map myeloid heterogeneity in its metabolic and cellular context. Cell Metab. 2022;34(8):1214–25 e6.

11. Kaya I, Michno W, Brinet D, Iacone Y, Zanni G, Blennow K, et al. Histology-Compatible MALDI Mass Spectrometry Based Imaging of Neuronal Lipids for Subsequent Immunofluorescent Staining. Anal Chem. 2017;89(8):4685–94.

12. Dufresne M, Guneysu D, Patterson NH, Marcinkiewicz MM, Regina A, Demeule M, et al. Multimodal detection of GM2 and GM3 lipid species in the brain of mucopolysaccharidosis type II mouse by serial imaging mass spectrometry and immunohistochemistry. Anal Bioanal Chem. 2017;409(5):1425–33.

13. Strittmatter N, Richards FM, Race AM, Ling S, Sutton D, Nilsson A, et al. Method To Visualize the Intratumor Distribution and Impact of Gemcitabine in Pancreatic Ductal Adenocarcinoma by Multimodal Imaging. Anal Chem. 2022;94(3):1795–803.

14. Yagnik G, Liu Z, Rothschild KJ, Lim MJ. Highly Multiplexed Immunohistochemical MALDI-MS Imaging of Biomarkers in Tissues. J Am Soc Mass Spectrom. 2021;32(4):977–88.

15. Lim MJ, Yagnik G, Henkel C, Frost SF, Bien T, Rothschild KJ. MALDI HiPLEX-IHC: multiomic and multimodal imaging of targeted intact proteins in tissues. Front Chem. 2023;11:1182404.

16. Rafols P, Torres S, Ramirez N, Del Castillo E, Yanes O, Brezmes J, et al. rMSI: an R package for MS imaging data handling and visualization. Bioinformatics. 2017;33(15):2427–8.

17. Rafols P, Heijs B, Del Castillo E, Yanes O, McDonnell LA, Brezmes J, et al. rMSIproc: an R package for mass spectrometry imaging data processing. Bioinformatics. 2020;36(11):3618–9.

