## Supplementary figures for "Integration of Mass Cytometry and Mass Spectrometry Imaging for Spatially Resolved Single Cell Metabolic Profiling"

**Supplementary data**

Supplementary table 1| IMC FFPE tumor immunophenotyping panel used for fresh frozen tissue after MALDI-MSI

| **Target** | **Clone** | **Metal** | **Incubation Time** | **Incubation Temp** | **Dilution** | **Company** |
| --- | --- | --- | --- | --- | --- | --- |
| CD4 | EPR6855 | 145 Nd | Indirect ON | RT | 100 | Abcam |
| CD8a | D8A8Y | 146 Nd | 5h | RT | 50 | Cell Signaling Technology |
| ICOS | D1K2T | 161 Dy | 5h | RT | 50 | Cell Signaling Technology |
| CD204 | J5HTR3 | 164 Dy | 5h | RT | 50 | Thermofisher Scientific |
| CD163 | D6U1J | 173 Yb | 5h | RT | 50 | Cell Signaling Technology |
| HLA-DR | TAL 1B5 | 141 Pr | 5h | RT | 100 | Abcam |
| CD11b | D6X1N | 144 Nd | 5h | RT | 100 | Cell Signaling Technology |
| Granzyme B | D6E9W | 150 Nd | 5h | RT | 100 | Cell Signaling Technology |
| CD14 | D7A2T | 163 Dy | 5h | RT | 100 | Cell Signaling Technology |
| CD7 | EPR4242 | 174 Yb | 5h | RT | 100 | Abcam |
| CD11c | EP1347Y | 176 Yb | 5h | RT | 100 | Abcam |
| CD45 | D9M8I | 149 Sm | Overnight | 4C | 50 | Cell Signaling Technology |
| CD3 | EP449E | 153 Eu | Overnight | 4C | 50 | Abcam |
| FOXP3 | D608R | 159 Tb | Overnight | 4C | 50 | Cell Signaling Technology |
| CD27 | EPR8569 | 175 Lu | Overnight | 4C | 50 | Abcam |
| Vimentin | D21H3 | 194 Pt | Overnight | 4C | 50 | Cell Signaling Technology |
| Keratin | C11 and AE1/AE3 | 198 Pt | Overnight | 4C | 50 | Biolegend/ CST |
| CD68 | D4B9C | 143 Nd | Overnight | 4C | 100 | Cell Signaling Technology |
| CD31 | 89C2 | 147 Sm | Overnight | 4C | 100 | Cell Signaling Technology |
| CD57 | HNK-1 / Leu-7 | 151 Eu | Overnight | 4C | 100 | Abcam |
| Ki-67 | 8D5 | 152 Sm | Overnight | 4C | 100 | Cell Signaling Technology |
| CD45RO | UCHL1 | 165 Ho | Overnight | 4C | 100 | Cell Signaling Technology |
| D2-40 | D2-40 | 166 Er | Overnight | 4C | 100 | Biolegend |
| CD38 | EPR4106 | 169 Tm | Overnight | 4C | 100 | Abcam |
| Histone H3 | D1H2 | 209Bi | Overnight | 4C | 50 | Cell Signaling Technology |

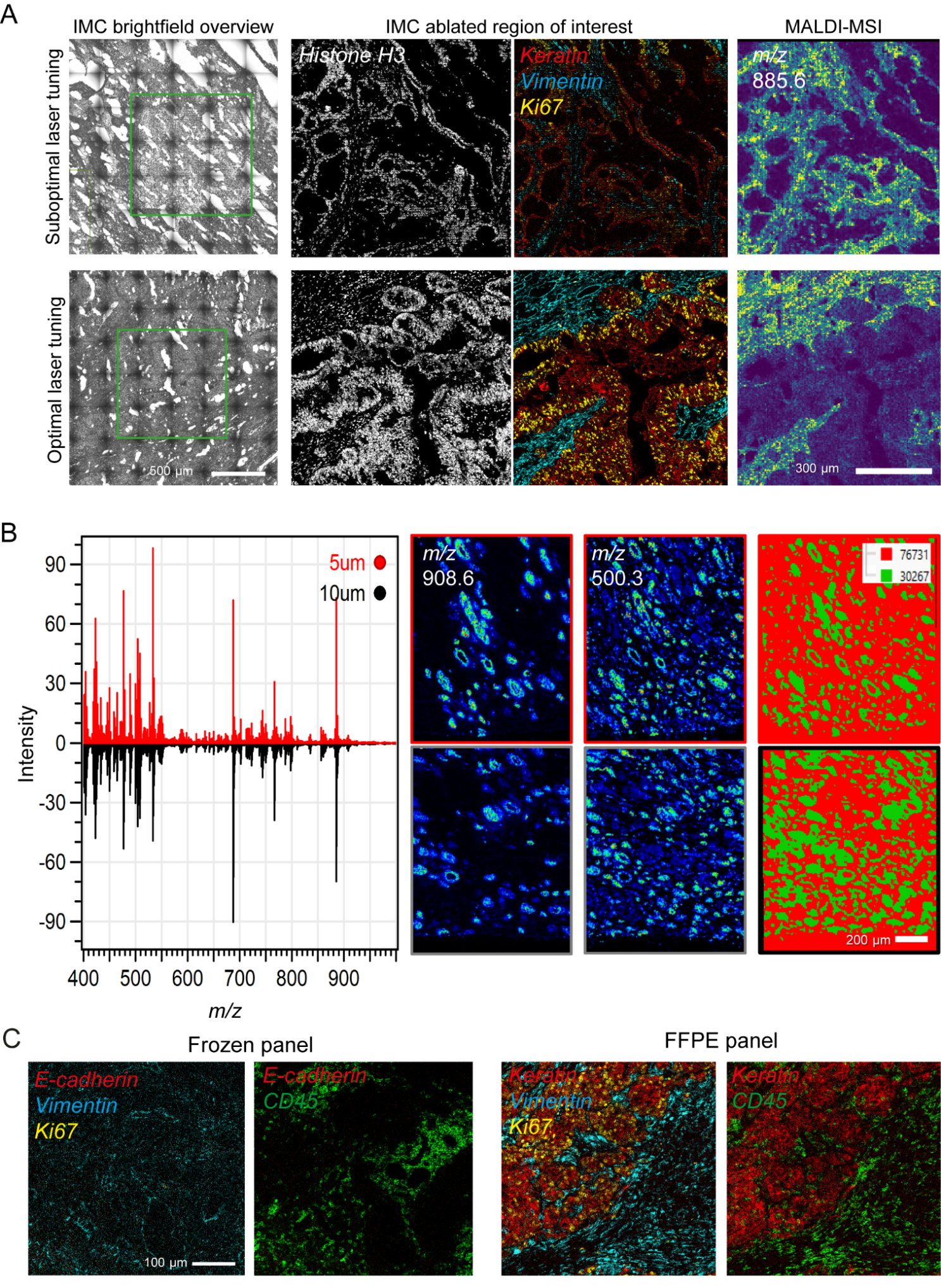

Supplementary figure 1| A: Left: brightfield overview-scan of tissue before IMC acquisition. The green square represents the selected region of interest (ROI). Middle: IMC ablated ROI with a single marker IMC image of histone and multi-marker IMC image. Right: non-normalized MALDI-MSI image of m/z 885.6 (assigned to PI(38:4) [M-H]^-^) shown in the viridis color scale. The top panel demonstrates the effect of a poorly adjusted MALDI laser, resulting in a grid pattern observed in the IMC brightfield overview and low IMC detection levels. The bottom panel shows the results after optimizing the MALDI laser. B: Comparison between total ion current (TIC) normalized average spectra of MALDI-MSI on a 5 µm (red) and 10 µm (black) thick tissue section of human kidney. TIC normalized peak intensity is represented by the viridis color scale. Spatial segmentation is performed using bisecting k-means clustering. C: Comparison of antibody panels and protocols optimized for fresh-frozen or FFPE tissues after MSI-IMC. Two images from the same ROI are shown for each antibody panel.

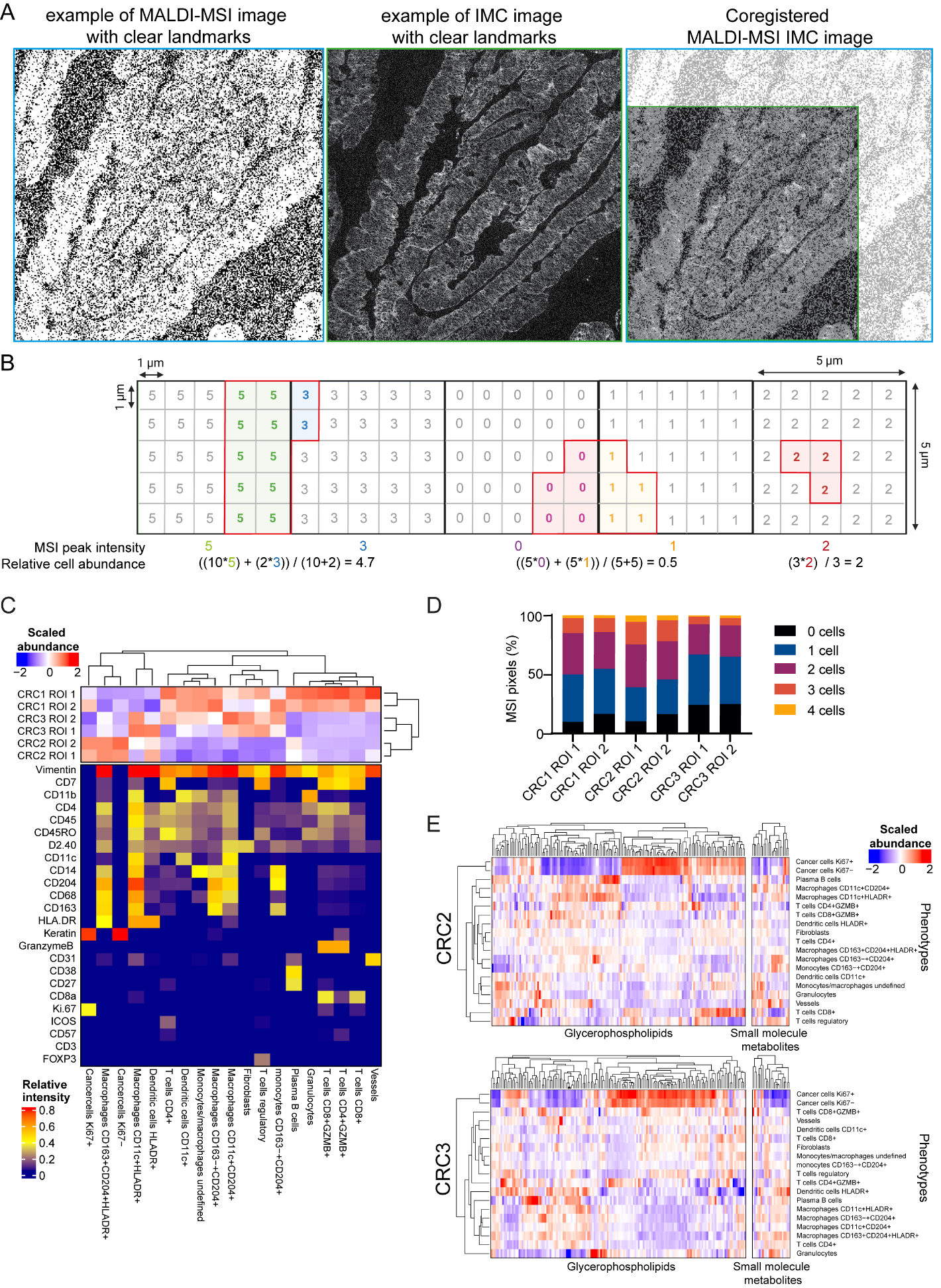

Supplementary figure 2| A. Approach for aligning MALDI-MSI and IMC data using visual landmarks present in both datasets. B. Schematic illustrating the approach for calculating metabolite abundance in each cell when pixel sizes differ between MALDI-MSI and IMC. Five 5x5 µm MALDI-MSI pixels are shown, each containing 25 1x1 µm IMC pixels. Cells within the pixels are coloured with a red border, and a cell can span multiple MALDI-MSI pixels. To calculate metabolite abundance per cell, the peak intensity of the overlapping 5x5 µm MALDI-MSI pixel was assigned to each 1×1 µm IMC-pixel. The IMC cell segmentation mask (red borders) and the assigned peak intensities of each 1x1 µm were combined, and the relative metabolite abundance per cell was calculated using the mean of all pixels within a cell. C. Relative marker expression and abundance per image of cellular phenotypes determined by IMC. D. distribution of cell numbers in a single MSI pixel for each image. E. Scaled metabolite expression profiles between IMC identified cellular phenotypes in CRC2 and CRC3. Hierarchical clustering was guided by glycerophospholipid abundances.

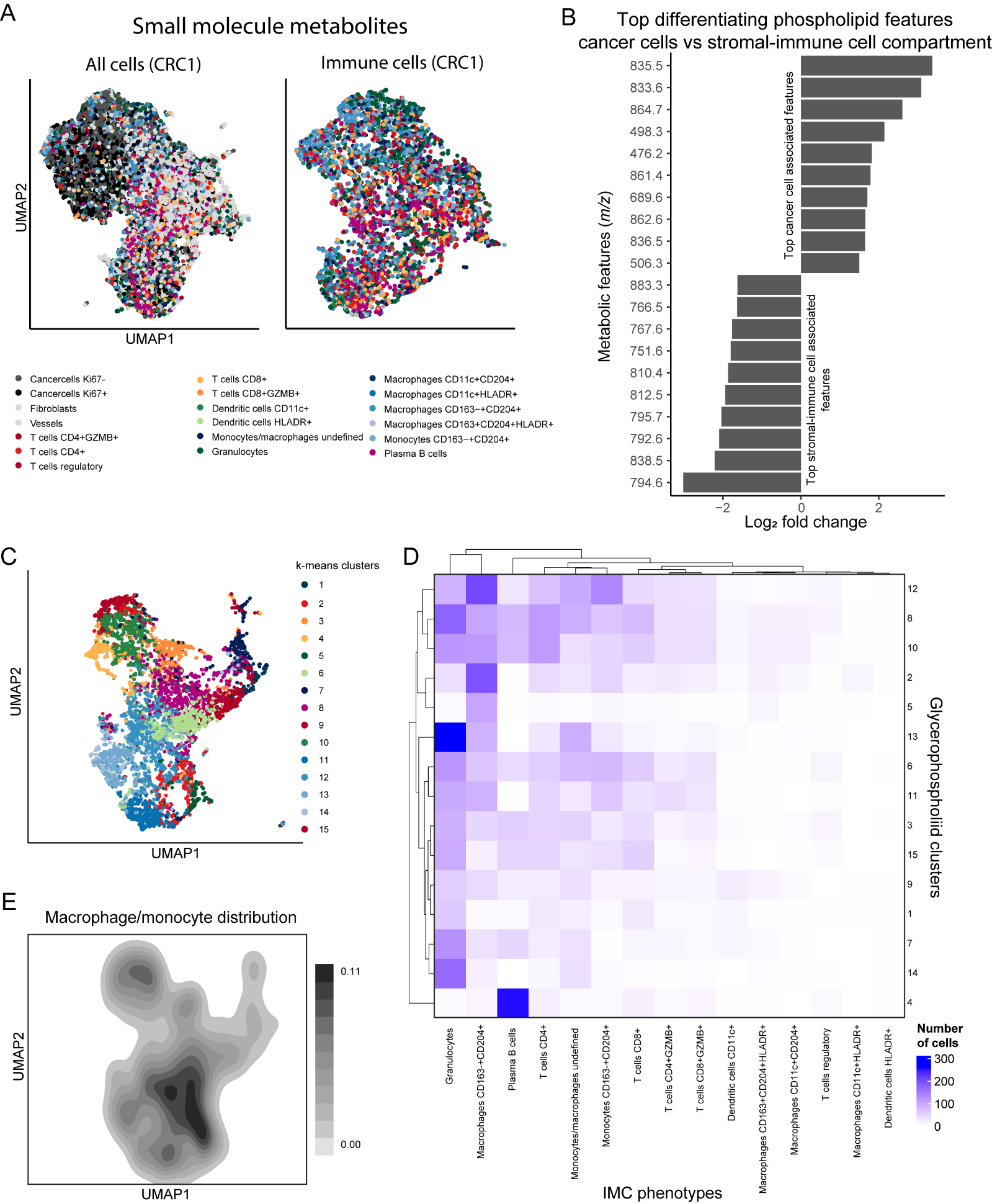

Supplementary figure 3| A. UMAP embedding of all cells or immune cells only of CRC1, clustered by small molecule metabolites (*m/z* <400). Cells were labelled according to their phenotypes. B. Top differentiating glycerophospholipid features between cancer cells and the stromal-immune cell compartment, calculated for all images in the dataset. C. k-means clustering of all immune cells in CRC1 based on glycerophospholipid features, visualized on the UMAP embedding shown in Figure 1C. D. Confusion matrix comparing IMC phenotypes with k-means clusters to determine clustering by cell type or glycerophospholipid feature. The heatmap indicates the number of cells that overlap between the two clustering methods.
